# Radical Footprinting in Mammalian Whole Blood

**DOI:** 10.1101/2024.09.29.615683

**Authors:** Mingming Zhao, Lyle Tobin, Sandeep K. Misra, Ajay Sharma, Juliette Locklar, Anter Shami, Sayed Mobarak, Haolin Luo, Lisa M. Jones, James A. Stewart, Joshua S. Sharp

## Abstract

Hydroxyl Radical Protein Footprinting (HRPF) is a powerful tool to probe protein higher-order structure, as well as protein-protein and protein-carbohydrate interactions. It is mostly performed *in vitro*, but recent advances have extended its use to live cells, nematodes, and 3D cultures. However, application in living mammalian tissues has not been accomplished. Here, we present the first successful use of radical protein footprinting (RPF) in mammalian whole blood from wild-type (WT) and type 2 diabetes mellitus (T2DM) BKS. Cg *Dock7*^*m*^ +/+ *Lepr*^*db*^/J mice. Using persulfate photoactivated with the FOX Photolysis System, we achieved effective protein labeling without significant disruption to blood cell morphology. An optimized quenching protocol eliminated background labeling. We report oxidative modifications in 11 selected proteins, revealing disease-associated conformational changes in multiple proteins. These findings demonstrate the feasibility of RPF in mammalian blood and open new opportunities for structural proteomics in preclinical models and clinical samples.

## INTRODUCTION

The biological function of protein is determined by higher order structure (HOS) ^1^. Characterizing protein HOS in their native environment is essential for understanding their function, but this remains challenging in complex mammalian biological samples. Hydroxyl radical protein footprinting (HRPF), as originally reported by Chance^2^ has become a powerful structural biology approach for elucidating protein HOS. Hydroxyl radicals (•OH) generated *in situ* diffuse to the protein surface, covalently modifying amino acid side chains at a rate proportional to their average solvent accessible surface area^3, 4^. The amount of oxidation at each site is measured by liquid chromatography coupled to tandem mass spectrometry (LC-MS/MS), generating a protein footprint. Alterations in the protein footprint under different conditions reflect changes in the protein HOS^5^. HRPF can probe protein dynamic conformational changes^6, 7, 8^, protein-protein and protein-ligand interactions^6, 7, 8^, and the conformational consequences of post-translational modifications^9^. Most recently, HRPF has been introduced to facilitate protein HOS analysis *in vivo* in living systems, including cancer spheroids^10^ and nematodes^11^.

Despite these advances, HRPF has not been demonstrated in any mammalian tissue, including blood, due to barriers like UV absorption by solid tissues. Another major technical hurdle to the implementation of HRPF in blood is the high concentration of catalase, which rapidly decomposes H_2_O_2_ (the predominant hydroxyl radical precursor for HRPF) before photolysis and •OH generation. For radical protein footprinting (RPF) to be achieved in blood, catalase must be inhibited, or an alternative radical precursor must be employed. The sulfate radical anion (SO_4_^−^•) was demonstrated for RPF by Gau and co-workers in 2010 and generates a protein footprint strikingly similar to that of •OH, possibly through a mixed sulfate and hydroxyl radical mechanism^12^. Although persulfate was recently employed for in-cell RPF in HEK cells^13^, it remains poorly characterized for use in RPF. No dosimetry method exists for SO_4_^−^• footprinting, which would detect and normalize the effective radical dose across replicates. Further, previous studies have photolyzed persulfate using custom excimer laser set-ups, which complicate standardization for clinical and many pre-clinical applications.

Here, we report the use of sodium persulfate for the first RPF of mammalian whole blood, using the biomedically relevant matrix of BKS. Cg *Dock7*^*m*^ +/+ *Lepr*^*db*^/J type 2 diabetes mellitus (T2DM; Homozygous for *Lepr*^*db*^; The Jackson Laboratory Strain 000642; db/db) mouse model as illustrated in **Figure 1**. We introduce the first SO_4_^−^• RPF with the Flash Oxidation (FOX) Photolysis System, a commercially available, standardized system that uses a broadband UV flash lamp^14^. We demonstrate both a real-time radical dosimetry system for *in vitro* SO_4_^−^• RPF as well as an effective secondary oxidant quench scheme that eliminates background oxidation from the RPF process. Using this workflow, we successfully performed RPF in whole murine blood, and oxidatively modified extracellular proteins. Comparison of protein footprint in non-diabetic, wild-type (WT) mouse (BKS.*Cg-Dock7m +/+ Leprdb/J*; Heterozygous for *Lepr*^*db*^, Heterozygous for *Dock7*^*m*^; Strain 000642) to our T2DM model murine blood revealed disease-associated protein conformational changes in two of the eleven proteins analyzed. Finally, we demonstrate that in-blood RPF accurately reflects protein topography by comparison with *in vitro* footprints of purified proteins. This work opens up powerful possibilities using RPF to investigate protein HOS in complex biological matrices, particularly whole blood, with potential applications in identifying molecular mechanisms underlying protein therapeutic efficacy, conformational biomarkers of disease, and novel drug targets, among myriad biological and pharmaceutical applications.

**Figure 1.**
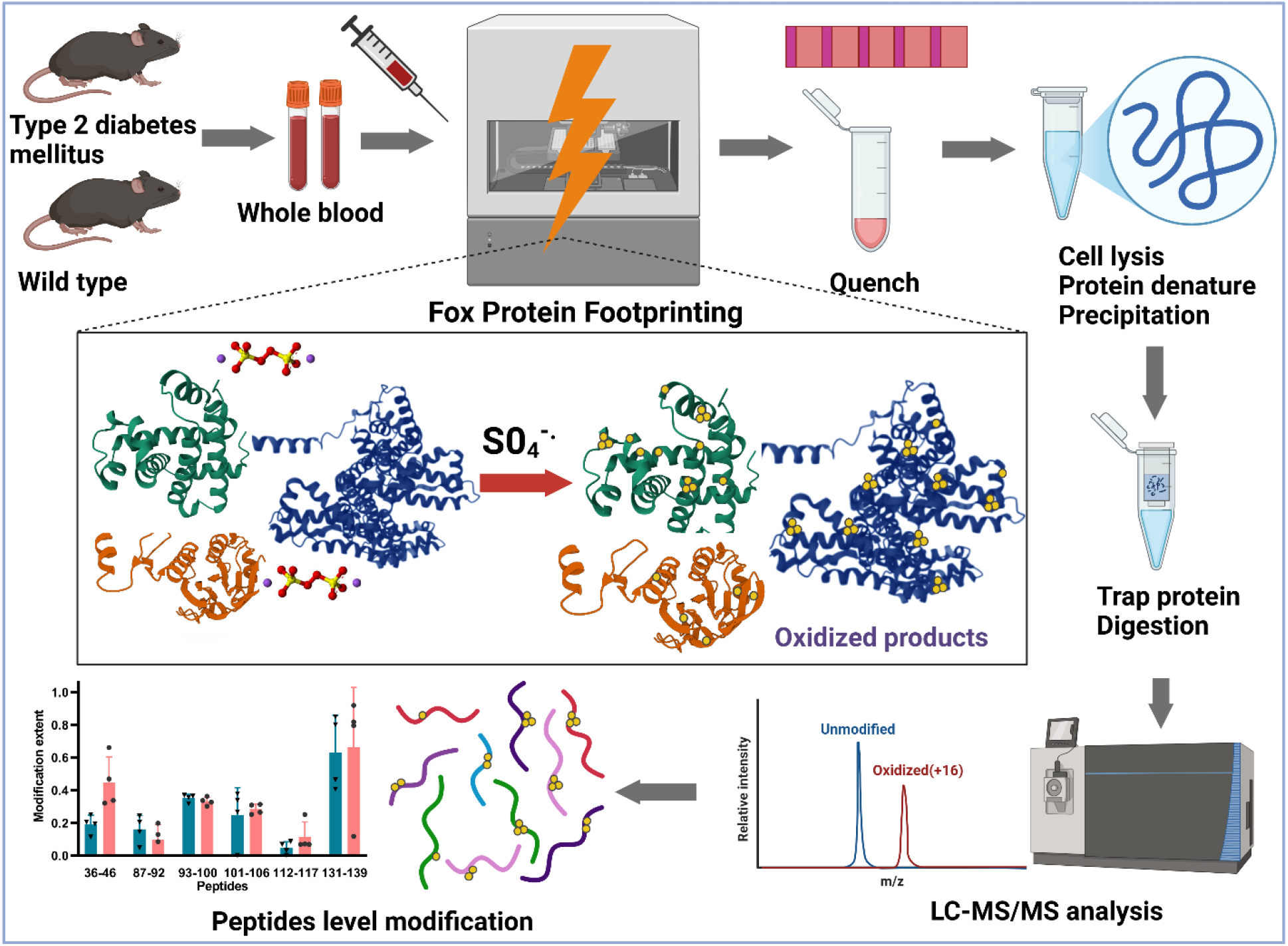
Experimental workflow of in-blood RPF. The figure was created with BioRender.com.

## MATERIALS AND METHODS

### Materials and Reagents

sTRAP micro columns were obtained from Protifi (Prod# CO2-micro-80, New York, NY, USA). Sodium chloride (NaCl), Sodium persulfate (Na_2_S_2_O_8_), equine myoglobin, bovine hemoglobin, N, N′-Dimethylthiourea (DMTU), iodoacetamide (IAA), hydroxylamine, sucrose, acetone, LC-MS grade of methanol, formic acid (FA), acetonitrile, water and isopropanol were obtained from Sigma Aldrich Corp. (St. Louis, MO). Triton X-100, imidazole, EDTA, and 13×100 mm test tubes were purchased from VWR International (Radner, PA). Adenine, 3-amino-1:2:4-triazole (3AT), dithiothreitol (DTT), Tris(2-carboxyethyl) phosphine (TCEP), sodium dodecyl sulfate (SDS), mono- and di-basic sodium phosphate, imidazole, aminoguanidine, 30% hydrogen peroxide (H_2_O_2_), MS-grade trypsin, EDTA-coated tubes (Catalog No.22-040-101), protease inhibitor tablets, and thiosemicarbazide (TSC) were purchased from Thermo Fisher Scientific (Waltham, MA). Methionine amide was purchased from Bachem (Bubendorf, Switzerland). Mouse complement c3 was acquired from Acro Biosystems (Newark, DE, USA). Bovine blood was generously provided by Home Place Pastures (Como, MS). Canine blood was generously provided by Dr. Harry Fyke, University of Mississippi. Initial wild-type mouse blood for method development was provided by Dr. Michael Bouvet at the University of California San Diego School of Medicine. Diabetic and wild-type mouse blood reported here was collected as described below in Animals and Whole Blood Collection.

### Assessment of Radical Precursor for In-Blood RPF using Foam Assay

Four previously reported catalase inhibitors (hydroxylamine, 3AT, aminoguanidine, and TSC) were assessed in blood via a test tube foam assay modified from Iwase *et. al*^*15*^. This assay correlates the height of trapped catalase-generated oxygen gas to catalase activity. Bovine blood was collected during routine exsanguination for beef production, immediately stabilized with 1.8 mg/mL EDTA, and flash frozen with dry ice/ethanol. 50 μL thawed blood was incubated with an inhibitor in a test tube, into which 50 μL of 2% Triton X-100 and 100 μL of 30% H_2_O_2_ or 2.5 M sodium persulfate were spiked. Foam height was recorded after 30 min. To optimize inhibition, we varied inhibitor concentrations, incubation duration, temperature, and volumetric (v/v) ratio of blood to inhibitor.

### Photoactivation of sodium persulfate using the FOX system

Samples containing 1 mM adenine and 50 mM sodium persulfate were irradiated by a FOX Photolysis system (GenNext Technologies, Inc., Half Moon Bay, CA) at 2 Hz. UV lamp voltage increased in 100 V increments from 600 to 900 V. Other triplicate samples were irradiated at 900 V and contained varied sodium persulfate concentrations (0, 50, 75, 100, 125, 150, 175 mM) or varied sucrose concentrations (0, 25, 50, 75, 100, 125 mM). Real-time changes in adenine absorbance were monitored at 265 nm (Abs_265_) by the inline dosimetry module^16^. Background oxidation was measured by injecting the above samples at 0 V and measuring oxidation post-quench.

### Sulfate radical quenching

The efficacy of the sodium persulfate quench solution was assessed by quantifying secondary oxidation products. To measure secondary oxidation, sodium persulfate was photoactivated and subsequently collected into quench solutions spiked with 5 μM myoglobin and 5 μM [Glu]1-fibrinopeptide B (GluB) to ensure that only oxidations occurring post-radical formation were measured. 20 μL samples containing 100 mM sodium persulfate, 1 mM adenine, and 25 mM sodium phosphate (pH 7.2) were injected into the FOX and irradiated at 900 V. Quench solutions consisted of 200 mM DMTU and 70 mM methionine amide, with or without 200 mM imidazole, in 25 mM sodium phosphate buffer (pH 7.2). The secondary oxidation extent was evaluated by quantifying the modified myoglobin and GluB.

### Blood cell morphology imaging

For blood cell morphology imaging, stabilized canine blood drawn during routine veterinary care was mixed 1:10 with either sodium persulfate or sodium chloride to a final concentration of 200 mM. The mixed blood was smeared on a microscope slide and imaged using a Zeiss Axio Imager M1 bright field microscope. Post-acquisition image manipulation was performed only to match background brightness and contrast.

### Animals and Whole Blood Collection

C57BL/6J (wild-type-WT; (*BKS*.*Cg-Dock7m +/+ Leprdb/J*; Heterozygous for *Lepr*^*db*^, Heterozygous for *Dock7*^*m*^; Strain 000642) and type 2 diabetes mellitus BKS. Cg *Dock7*^*m*^ +/+ *Lepr*^*db*^/J type 2 diabetes mellitus (db/db; Homozygous for *Lepr*^*db*^; The Jackson Laboratory Strain 000642) mice were obtained and housed at the i3S animal facility under specific pathogen-free conditions. Animals were group-housed in an AAALAC-approved animal facility following the National Institutes of Health “Guide for the Care and Use of Laboratory Animals.” Mice experienced a 12h/12h light/dark cycle, and food (2014S Teklad Diets) and water were *ad libitum*. The University of Mississippi Animal Care and Use Committee (IACUC protocol number 23-011) approved all animal usage protocols.

At 16 weeks of age, the animals were weighed, and their blood glucose levels were measured (**Supplementary Table S1**). Mice were euthanized by CO_2_ asphyxiation followed by cervical dislocation to collect whole blood, which was drawn into EDTA-coated tubes and maintained at room temperature during transport. Blood samples were irradiated within 4 hours of collection.

### In-Blood SO_4_^−^• RPF

EDTA-stabilized whole blood was collected respectively from four db/db mice and four WT control mice. In-Blood RPF was performed for each mouse blood draw using a FOX Protein Footprinting System (GenNext Technologies, Inc., Half Moon Bay, CA). The flow rate was maintained at 15 μL /min to the desired FOX exclusion volume. For each injection, 20 μL samples were injected to overload the FOX Photolysis System’s injection loop. The FOX Photolysis System’s fluidics module pumps the sample using phosphate buffer saline as the carrier fluid. Immediately prior to injection, 2 μL of sodium persulfate or sodium chloride at final concentration of 200 mM was added to 18 μL whole blood sample. The whole blood of all biological samples was irradiated in technical duplicate under three experimental conditions: sodium persulfate and 0 V (unilluminated control); sodium persulfate and 800 V (RPF); sodium chloride at 0 V (negative control). Immediately after labeling, whole blood samples were collected at 20 μL of quench solution containing 200 mM imidazole, 200 mM DMTU and 70 mM methionine amide.

### Whole blood proteome sample preparation

After RPF and quench, samples were precipitated with four volumes of cold acetone and stored at –20 °C overnight. Thawed samples were centrifuged (14,000 × g, 10 min), washed twice with PBS, and resuspended in 300 μL of 50 mM Tris buffer. Proteins were lysed and digested using Profiti Micro S-Trap columns (CO2-micro-80) with minor modifications. Samples were lysed in 10% SDS buffer, ultrasonicated (5 min, 50 N, 1 kHz), reduced with 5 mM TCEP at 65 °C for 20 min, and alkylated with 20 mM IAA in the dark for 40 min. Proteins were trapped, washed, digested overnight with trypsin (1:10 ratio), and sequentially eluted with Tris buffer, formic acid, and acetonitrile.

### *In vitro* SO_4_^−^• RPF of complement c3

Protein labeling samples in a total volume of 20 μL consisting of 1 mM adenine, 25 mM sodium phosphate, pH 7.2, and 200 mM sodium persulfate were irradiated at 800 V and collected in the same quench solution as used in blood. To digest the protein samples, a final concentration of 100 mM Tris, pH 8.0, and 1 mM sodium chloride were added to the samples and heated at 95 ^0^C for 15 min and immediately placed on ice for 2 min. 333 ng trypsin was added to the samples and incubated at 37 ^0^C with rotation for 14 hours. A final concentration of 0.1% formic acid was added to the samples to stop trypsin activity.

### Liquid chromatography mass spectrometry (LC-MS) analysis of eluted peptides

Eluted peptides were analyzed by an Orbitrap Fusion Tribrid mass spectrometer coupled with a Dionex Ultimate 3000 system nanoLC system (Thermo Fisher, CA). Samples were loaded onto a trap column (Thermo Fisher C18 5 *μ*M, 0.3 × 5 mm) and peptide separation was achieved through a nano C18 column (Acclaim 2 *μ*M, 0.075 × 150 mm) at a flow rate of 0.3 *μ*L/min. Solvent A was 0.1% formic acid in water, and solvent B was 0.1% formic acid in acetonitrile. The gradient consists of 2% solvent B held for 2 min, 2–7% solvent B over 2 min, 7–32% solvent B over 58 min, 32–95% solvent B over 3min and held for 3 min, returned to 2% solvent B over 4 min and held for 8 min. The total gradient run time was 80 min.

Mass spectrometric analysis was performed on an Orbitrap Fusion Tribrid in data dependent acquisition mode using Xcalibur software (Thermo-Fisher Scientific, Waltham, MA). Mass spectra of the peptides were analyzed over a 250–2000 *m*/*z* range at normal mass resolving power with a dynamic exclusion (Orbitrap Resolution 60000). Peptides with single charge state were excluded. Electrospray ionization was carried out in a positive ion mode. The ion source voltage was kept at 2450V. Data-dependent mode MS/MS was used, with CID applied to each selected precursor. The CID intensity threshold was set at 5.0 × 10^3^, and the CID collision energy was set at 32% with a rapid ion trap scan.

### Proteomic data processing

Byonic (v4.4.2, Protein Metrics, San Carlos, CA) was used to identify the peptide and determine the protein sequence coverage. LC-MS data were searched against the complete *Mus musculus* Proteome. Trypsin cleavage specificity was set to Lys, and Arg residues, except before Pro residues, with up to 2 missed cleavages permitted. Precursor mass tolerance was set to 10 ppm, and the fragment mass tolerance was set to 0.4 Da. Met oxidation was set as a variable modification and Cys residues were set for carbamidomethylation modification. Protein false discovery rate was set at 1%. The identified proteins in whole blood were ranked by average sequence coverage to select the top eleven proteins for footprinting analysis. Peptide-level oxidation was quantified using Foxware v2.1.2 (GenNext Technologies, CA).

### Statistical analysis

Each data point represents a biological replicate that is the mean of two technical replicates. All data are expressed as the mean ± standard deviation (SD). A two-tailed unpaired t-test was used to compare two groups. One-way analysis of variance (ANOVA) followed by Dunnett’s test was used to compare three or more groups (P < 0.05 was considered statistically significant difference, NS means not significant). GraphPad Prism was used for analysis.

## RESULTS

### Selection of a stable radical precursor for whole blood

Most previous cell-based and *in vivo* radical footprinting experiments have used hydrogen peroxide-based FPOP for fast radical generation^10, 17^. However, blood notably displays high catalase activity^18^, which complicates the use of hydrogen peroxide as a radical precursor. We tested four catalase inhibitors to determine if catalase and/or peroxidase activity in whole blood could be sufficiently reduced without altering the gross physiological properties of the blood sample. We also assessed gas generation in blood from sodium persulfate, previously utilized for in-cell FPOP^13^. Of the four inhibitors, hydroxylamine was the most effective, with a 95% reduction in catalase activity at 200 mM concentration, 5:1 v/v blood to inhibitor stock, and room temperature as seen in **Figure 2 A**. Longer incubation and reduced temperature improved inhibition (data not shown).

**Figure 2.**
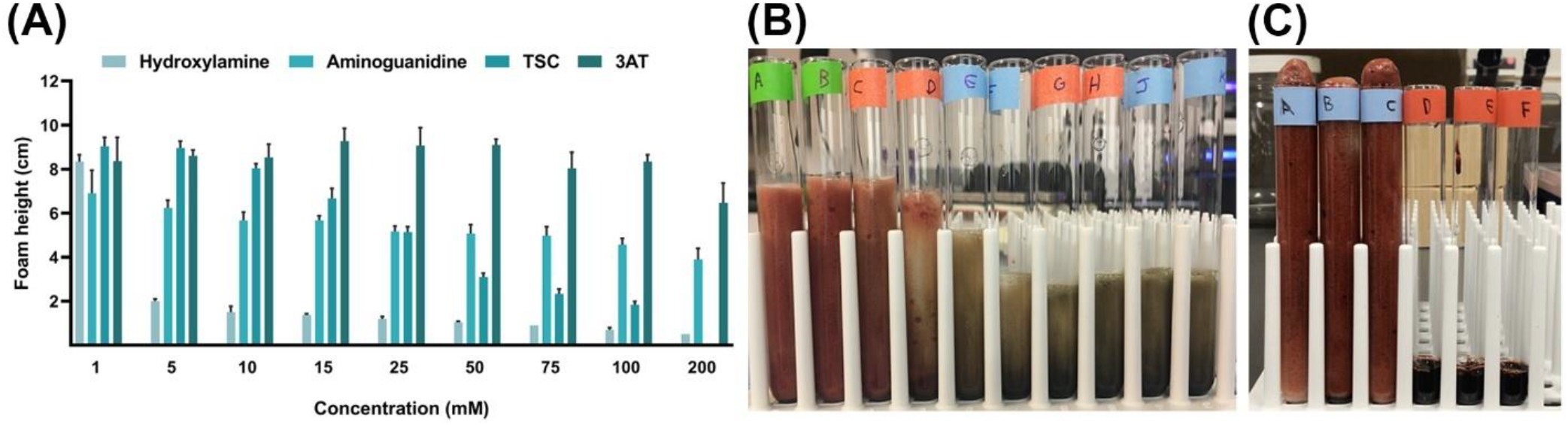
Test of inhibitors of catalase. **(A)** Foam height after 30 minute incubation (5:1 v/v blood to inhibitor) at room temperature. TSC is insoluble at 200 mM. Error bars represent one standard deviation of triplicate measurement. **(B)** Representative samples of blood spiked with varying final concentrations of hydroxylamine: 0 mM **(green A-B)**, 5 mM **(orange C-D)**, 25 mM **(blue E-F)**, 100 mM **(orange G-H)**, and 142 mM **(blue J-K). (C)** Blood spiked with equal volumes of 30% H_2_O_2_ **(blue)** or 2.5 M sodium persulfate **(red)**.

However, TSC and hydroxylamine induced gross changes in blood coloration and constitution, clearly disrupting the native blood environment. At a more physiologically relevant 20:1 v/v, substantial catalase activity remained. Importantly, none of the four inhibitors prevented foam generation in the blood at concentrations compatible with physiological conditions. Barring the identification of some other effective inhibitor, HRPF is likely not feasible in whole blood with H_2_O_2_ as a radical precursor. Conversely, when sodium persulfate was spiked into blood at final 200 mM concentration as shown in **Figure 2B-C**, no gas generation or gross changes in blood properties were observed. Therefore, sodium persulfate was used for all in-blood RPF experiments.

### Photoactivation of sodium persulfate using the FOX system

While RPF has been reported using sodium persulfate with a KrF excimer laser^12, 13^, sodium persulfate has not been documented as an effective radical precursor using the FOX Footprinting system. We tested whether persulfate could be effectively photoactivated by the FOX flash lamp and whether the inline adenine dosimetry was compatible with SO_4_^−^• RPF *in vitro*. Dosimetry results are presented in **Figure 3**. When adenine is oxidized, the loss of UV absorbance signal at 265 nm allows effective measurement of the radical generation and scavenging. As sulfate and hydroxyl radicals are broadly reactive and can be consumed by solutes like buffers and excipients^19^, real-time dosimetry allows for compensation to ensure that analytes receive equivalent radical doses^20, 21, 22^. Accordingly, we examined the relationship between in-line UV absorbance of 1 mM adenine and sulfate radical effective dose under various conditions.

**Figure 3.**
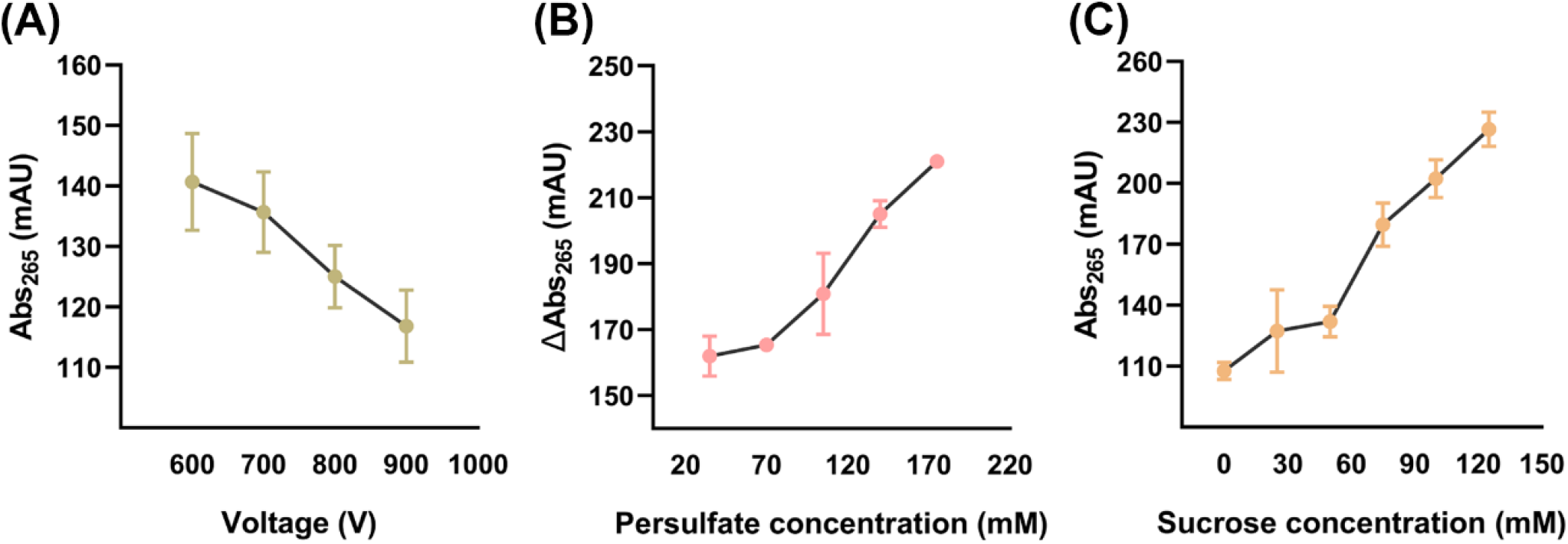
Adenine dosimetry response to SO_4_^−^•. **(A)** Adenine absorbance after illumination using different lamp voltages, **(B)** change in absorbance for different persulfate concentrations at 900 V versus 0 V, and **(C)** adenine absorbance at 900V in the presence of different concentrations of sucrose radical scavenger. Error bars represent one standard deviation of a triplicate measure.

Adenine dosimeter response was assessed first in response to an increase in the effective radical dose, achieved via increased light fluence as modulated by flash voltage. Abs_265_ decreased from ∼140 mAU to ∼116 mAU as lamp voltage was increased from 600V to 900V in the presence of 50 mM persulfate (**Figure 3A**), indicating an inverse relationship between adenine absorbance and effective radical dose, similar to that observed with hydroxyl radicals. Adenine dosimetry obtained with varied radical precursor concentrations, however, is confounded by the intrinsic Abs_265_ of sodium persulfate. Thus, Abs_265_ was recorded for triplicate samples containing 1 mM adenine and varied persulfate concentrations that were irradiated at either 0V or 900V. These values were compared to find the magnitude of the decrease in Abs_265_, hereafter denoted by ΔAbs_265_. As seen in **Figure 3B**, ΔAbs_265_ increased by 0.45 mAU per mM persulfate again indicating an inverse relationship between adenine absorbance and effective radical dose once corrected for the absorbance of the persulfate precursor. To determine that dosimetry is sensitive to radical scavenging rather than just a measurement of radical generation, concentration of a radical scavenger (sucrose) was increased across triplicate samples to decrease effective radical dose. This yielded a linear response within the range tested, with Abs_265_ increasing by 0.992 mAU per mM sucrose (**Figure 3C**), again verifying that adenine absorbance varies inversely with effective radical dose.

These dosimetry results show that sodium persulfate can be photoactivated using the FOX Foot-printing system, and that adenine can be a highly reliable SO_4_^−^• dosimeter in the appropriate buffer systems if the persulfate concentration is kept constant. This allows for the real-time compensation of *in vitro* conditions across SO_4_^−^• RPF experiments. However, compound changes of persulfate concentration and either lamp voltage or scavenger concentration must carefully account for the inherent Abs_265_ of persulfate. We recommend only changing lamp voltage for compensation efforts^20, 21^ in persulfate-based labeling when possible.

### Selection of an effective quench solution for SO_4_^−^•

Recent observations suggest that standard HRPF quenching protocols are not sufficient for SO_4_^−^• RPF (L.M. Jones, unpublished results), contrary to previous reports^12, 13^. An effective quench is critical as secondary oxidation after primary radical modification can introduce artifactual modifications to non-native structures^23^. In this study, two quench solution systems were tested. Quench solution ? contained 200 mM DMTU and 70 mM Met amide − a standard HRPF quench that has been demonstrated effective for photoactivated H_2_O_2_^24^. Quench solution ? included 200 mM imidazole in addition to 200 mM DMTU and 70 mM methionine amide. The efficacy of the sodium persulfate quench solutions was assessed by quantifying secondary oxidation products. To induce secondary oxidation, sodium persulfate was photoactivated and subsequently collected into quench solutions already containing myoglobin and GluB. Quench efficacy was assessed at flash lamp voltages of 900V and 0V. In the 0V condition, myoglobin or GluB was injected into the FOX with persulfate before collecting in a quench to test background oxidation and the ability of an effective quench to limit it.

The results of these quenching experiments are shown in **Figure 4** and **Supplementary Figure S1**. We detected seven peptides from myoglobin. During electrospray ionization, significant insource oxidation occurred for peptide 2–17, even in samples without persulfate. This in-source oxidation can be differentiated from pre-existing oxidation by the perfect chromatographic co-elution of the in-source oxidation product with the unmodified peptide. These in-source oxidation products are consistent in intensity and can be observed in the no-persulfate controls. Some peptides exhibited a significant increase in background oxidation in quench solution I which only contains DMTU and methionine amide. For instance, peptide 120–134 showed a three- to four-fold increase in modification in quenches lacking imidazole. Quenches with 200 mM imidazole showed no significant difference in peptide modification with or without sodium persulfate. Moreover, there was not a significant increase in oxidation when myoglobin was injected in-line with persulfate without illumination as reported by Gau *et al*.^12^. These results indicate that a quench solution containing DMTU, methionine amide and imidazole effectively prevents background secondary oxidation artifacts in SO_4_^−^• RPF experiments on the FOX system.

**Figure 4.**
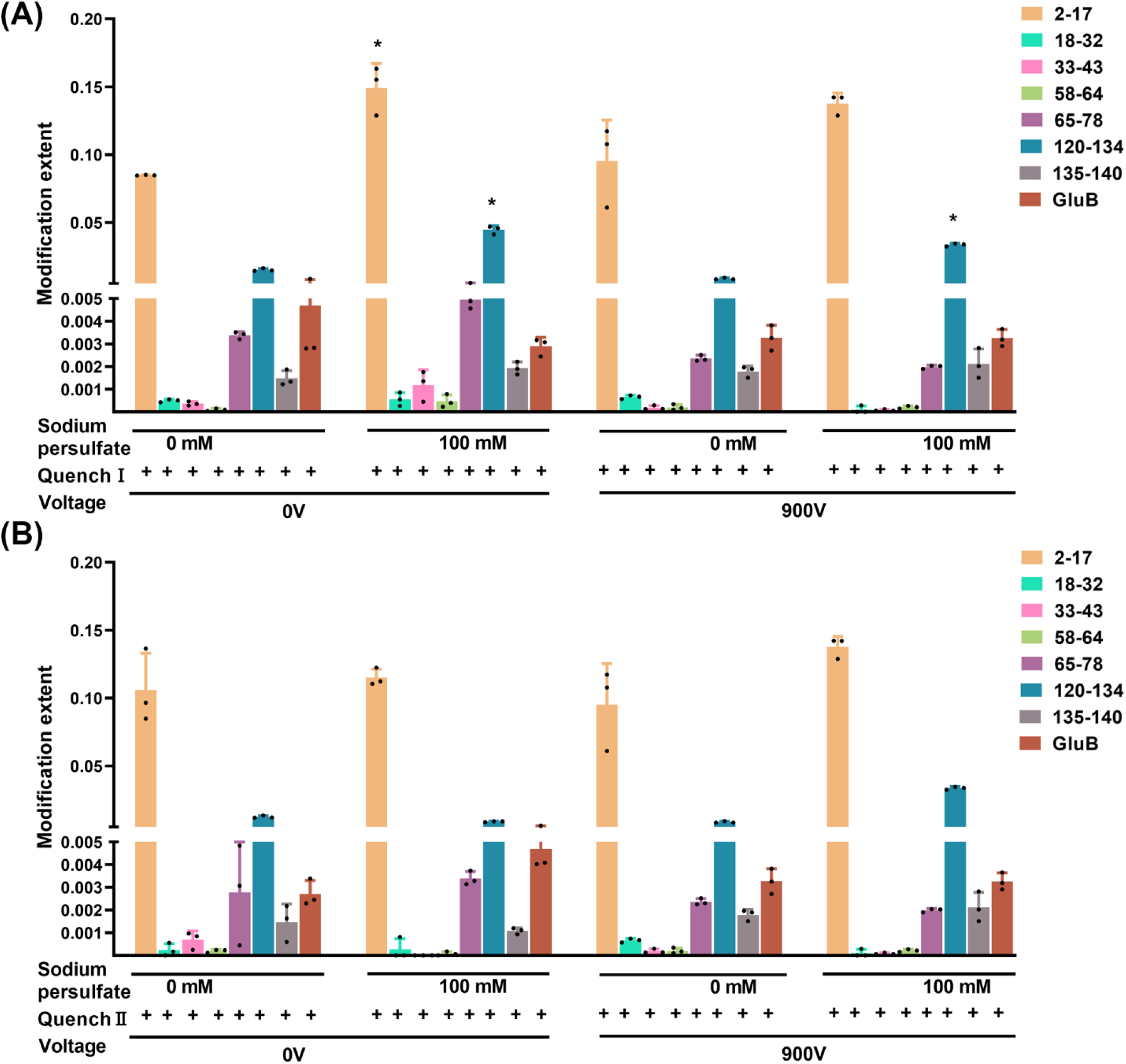
Selection of effective quench solutions for background oxidation. **(A) Evaluation of Quench I (without imidazole); (B) Evaluation of Quench ? (with imidazole)**. Statistical significance was calculated by a two-tailed unpaired t-test (α = 0.05) versus the 0 mM persulfate control under the same conditions; peptides showing a significant difference in oxidation are marked with an asterisk. Myoglobin peptides are denoted by their position. The myoglobin peptide corresponding to the 2–17 residues displays significant in-source oxidation. Error bars represent one standard deviation.

### Effect of sodium persulfate on blood cell morphology

To determine if SO_4_^−^• RPF preserved native-like conditions in stabilized whole blood before photoactivation, canine blood was exposed to either sodium persulfate (200 mM final concentration) or sodium chloride (200 mM final concentration). Cell morphology was examined using bright-field microscopy. Results are shown in **Figure 5**. 200 mM sodium persulfate induces modest morphological changes consistent with mild hypertonicity. Notably, an equivalent concentration of sodium chloride shows comparable, if not slightly more pronounced, changes in cell morphology. No other morphological change or evidence of increased cell lysis was observed, suggesting that the only gross morphology change from persulfate treatment is mild hypertonicity.

**Figure 5.**
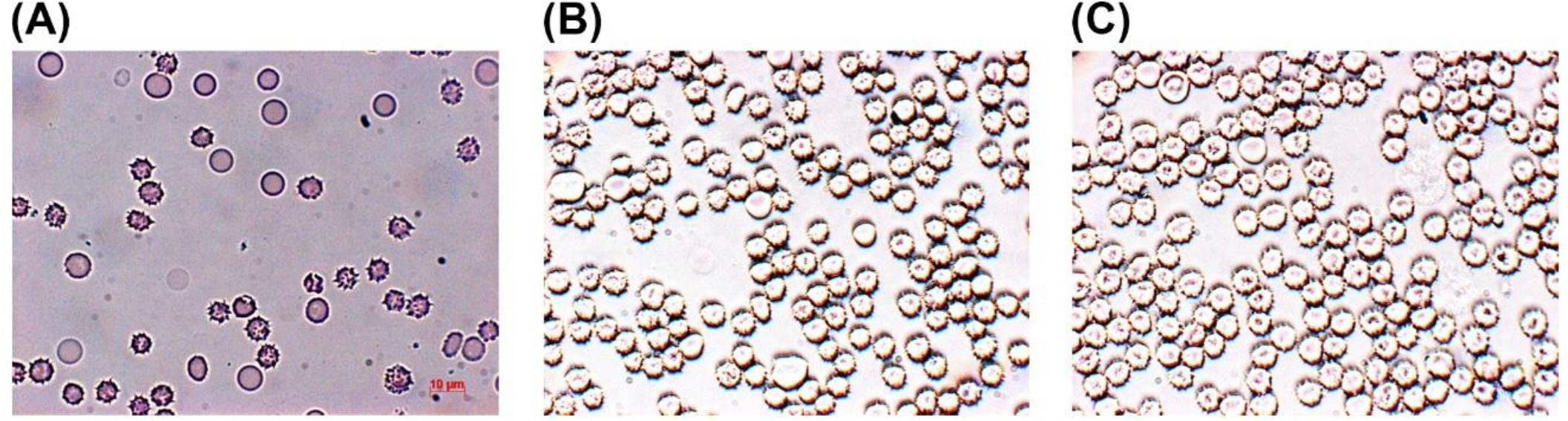
Effect of sodium persulfate on blood cell morphology. **(A)** Blood cells from canine blood. **(B)** Same blood with 200 mM sodium persulfate. Cells show a moderate increase in hypertonicity, evidenced by an increase in wrinkled cells. **(C)** An equal concentration of sodium chloride results in cells with an identical morphology to the persulfate sample

### Peptide-level modification of intracellular proteins in whole blood by SO_4_^−^• RPF

The resulting modified murine blood was subjected to a bottom-up proteomics workflow. To determine whether the blood proteome was successfully modified, we focused on the top 11 proteins identified by Byonic with highest sequence coverage. (The average sequence coverage of each of these proteins was higher than 50%, as shown in **Supplementary Table S2**). Oxidation was quantified at the peptide level. Interestingly, significant differences in modification extent were found between extracellular and intracellular proteins. As shown in **Figure 6 A-B**, in both WT and T2DM mouse whole blood samples, significantly higher modification coverage was observed in extracellular proteins compared to intracellular proteins. As demonstrated in **Supplementary Figure S2**, in WT mice whole blood, 54 peptides derived from extracellular proteins were detected, of which 20.0% exhibited an average modification extent (AME) per peptide greater than 0.3. In contrast, 68 peptides from intracellular proteins were identified, with only 1.5% showing an AME above 0.3. Similarly, in T2DM mouse whole blood, 13% of extracellular peptides were oxidatively modified to an extent greater than 0.3, while no intracellular peptides exhibited a modification extent above this threshold.

**Figure 6.**
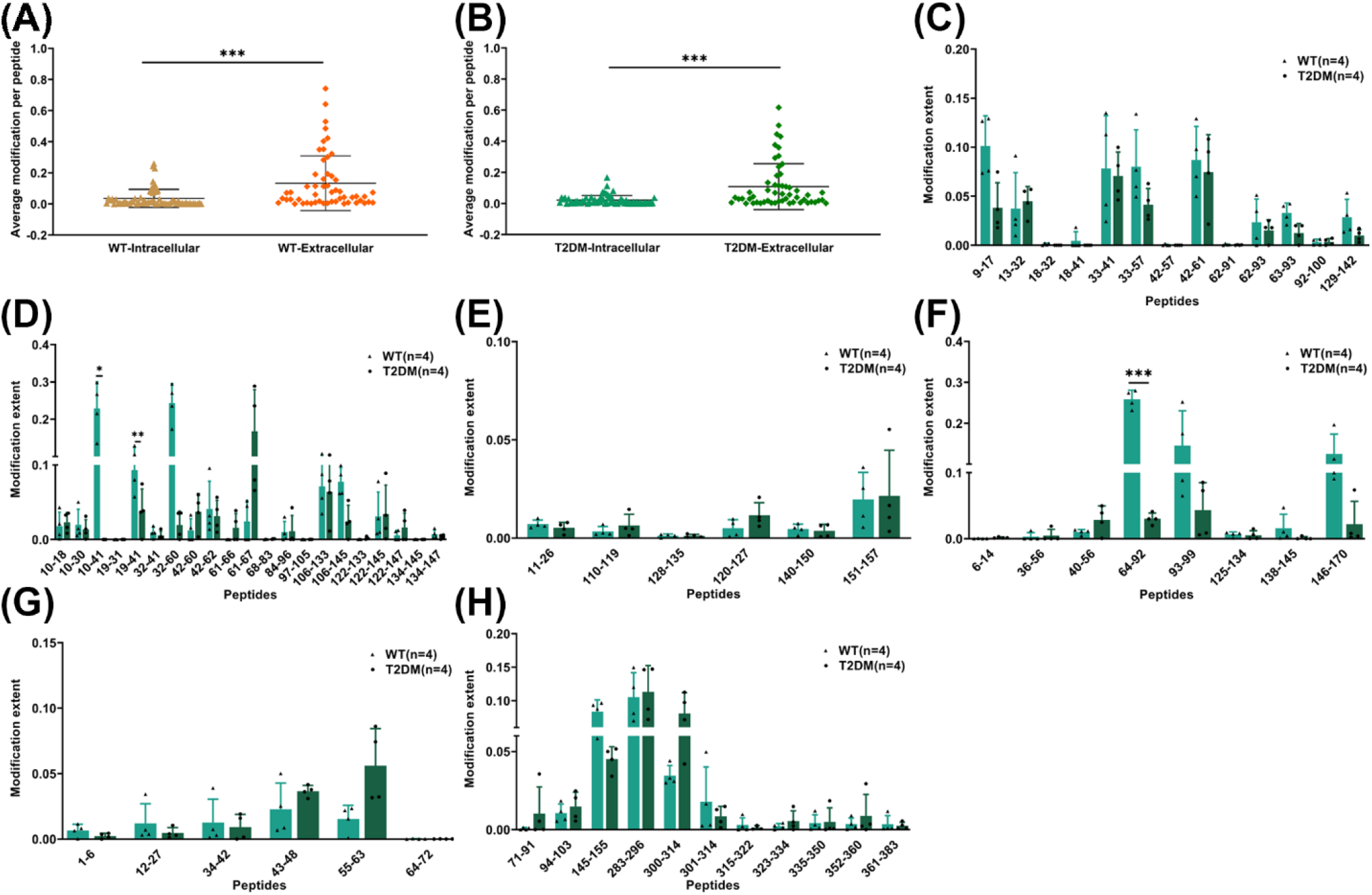
Comparison of in-blood RPF of top 6 intracellular proteins detected in WT and T2DM whole blood. Significantly higher modification was observed in extracellular proteins of WT **(A)** and T2DM **(B)** compared to intra-cellular proteins, Each data point represents a peptide.The modification extent at 0V was subtracted for each peptide; **(C-H)** Peptide-level oxidation of peptides from six intracellular proteins, after subtraction of 0V background oxidation values: **(C)** alpha-globin; **(D)** beta-globin; **(E)** peroxiredoxin-2**; (F)** Flavin reductase**; (G)** 60S ribosomal protein-L40; **(H)** Serine protease inhibitor. * p < 0.05, ** p < 0.01, *** p < 0.001 by unpaired T-test after Šídák correction.

To determine whether the observed discrepancies were due to differences in the abundance of intracellular versus extracellular proteins, we performed hierarchical clustering heatmaps based on the intensity of proteins detected by Byonic. As shown in **Supplementary Figure S3**, alpha and beta globin—both intracellular proteins—exhibited the highest abundance across all T2DM and WT samples. Overall, intracellular proteins displayed comparable, if not higher, abundance than extracellular proteins. These results suggest that the reduced extent of oxidation among intracellular proteins is not due to lower abundance. Rather, it is more likely attributed to the limited cell permeability of sodium persulfate. This indicates that SO_4_^−^• RPF can be successfully performed in mammalian whole blood and is capable of substantially modifying extracellular proteins. Efficient modification of intracellular proteins may require additional steps to increase reagent permeability^25^.

Additionally, as illustrated in **Figure 6 C-H**, we subtracted the background oxidation (0 V) and localized the peptide-level modifications in the top six intracellular proteins. Three peptides showed significant differences in the extent of modification between WT and T2DM mice after Šídák correction for the number of oxidized peptides measured in the protein. These included peptides 10–41 and 30–60 from beta-globin, and peptide 64–92 from flavin reductase.

### Peptide-level modification of extracellular proteins in whole blood by SO_4_^−^• RPF

The increased modification coverage in extracellular proteins provided greater structural information. As demonstrated in **Figure 7 A–E**, a total of 54 peptides from the top five extracellular proteins were modified. For each protein, at least one peptide exhibited a modification extent greater than 0.3. Oxidation coverage of the detected peptides ranged from sparse to very high, which is consistent with reported findings^26,27^, suggesting that optimization for protein system(s) of interest may permit interrogation of any target of interest in the blood proteome.

**Figure 7.**
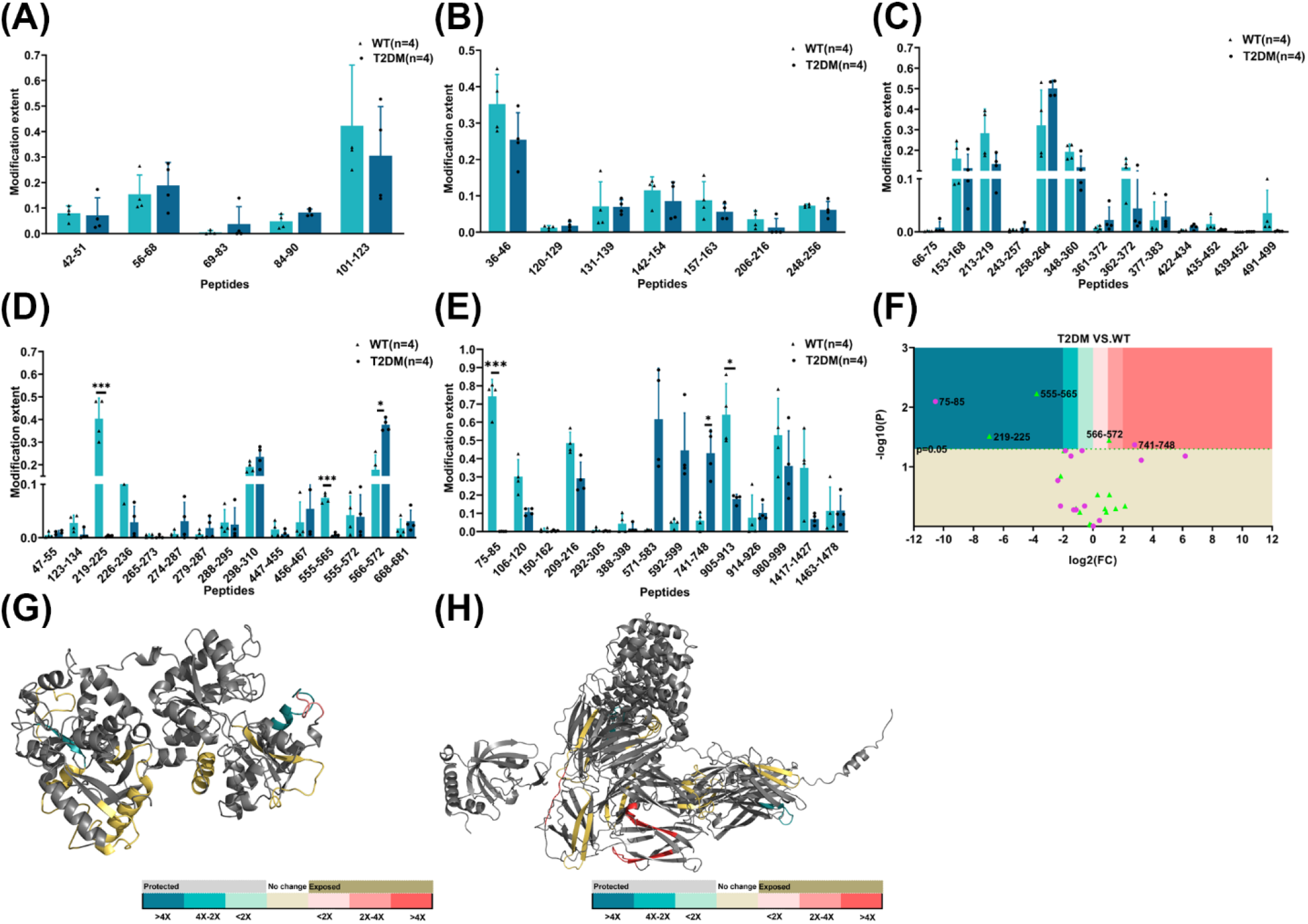
Comparison of the in-blood RPF of top extracellular proteins detected in WT and T2DM whole blood. **(A)** Transthyretin; **(B)** Apolipoprotein A-I; **(C)** Albumin; **(D)** Serotransferrin; **(E)** Complement C3; **(F)** Volcano plot of disease-associated conformational changes in T2DM, pink dots: peptides from complement c3, green dots: peptides from serotransferrin, **(G)** Topographical changes of protein serotransferrin in T2DM: peptide color represents fold change in oxidation, colors coded to match with **(F)**, gray peptides showed no measurable oxidation; **(H)** Topographical changes of protein complement c3 in T2DM: peptide color represents fold change in oxidation, colors coded to match with **(F)**, gray peptides showed no measurable oxidation. The modification extent at 0V was subtracted for each peptide. All statistics performed were unpaired two-tailed t-test with Šídák correction; * p < 0.05, ** p < 0.01, *** p < 0.001 by unpaired t-test.

The highest levels and broadest modification coverage were observed in two proteins: complement c3 and serotransferrin. To further interrogate disease-associated protein conformational changes, we probed the peptide-level modification data to identify in-blood SO_4_^−^• RPF oxidation patterns in WT and T2DM environment. These results were mapped onto the crystal structures of each protein **(Figure 7 F-H)**. Peptide 75–85 from protein complement c3 was significantly protected from oxidation in T2DM, compared to WT after Šídák correction. Notably, it showed an average modification extent of 0.74 in WT samples, whereas it remained unmodified in T2DM samples. In contrast, peptides 571–583 was significantly exposed to oxidation in T2DM whole blood, compared to WT. In the case of protein serotransferrin, two peptides, including 219 – 225, 555–565, were significantly protected from oxidation in T2DM, while one peptide, 566–572 was significantly exposed to oxidation in T2DM. These findings suggest that distinct, disease-associated alterations in protein surface solvent accessibility are detectable by in-blood RPF.

### Testing the influence of HOS on in-blood RPF

To determine whether the oxidation pattern observed in whole blood was reflective of the protein’s native structure, we corelated the footprint of complement c3 from murine whole blood with our *in vitro* footprint of mouse complement c3 (95% sequence identity, 90% sequence similarity). The correlation between *in vitro* and in-blood results is shown in **Figure 8**. Because the effective radical dose between *in vitro* and in-blood cannot be normalized due to a lack of effective radical dosimetry for in-blood measurements, we measured the Spearman correlation between the two experiments, finding a strong correlation (r=0.8286). This analysis supports that in-blood RPF reasonably reflects the topography of the measured proteins.

**Figure 8.**
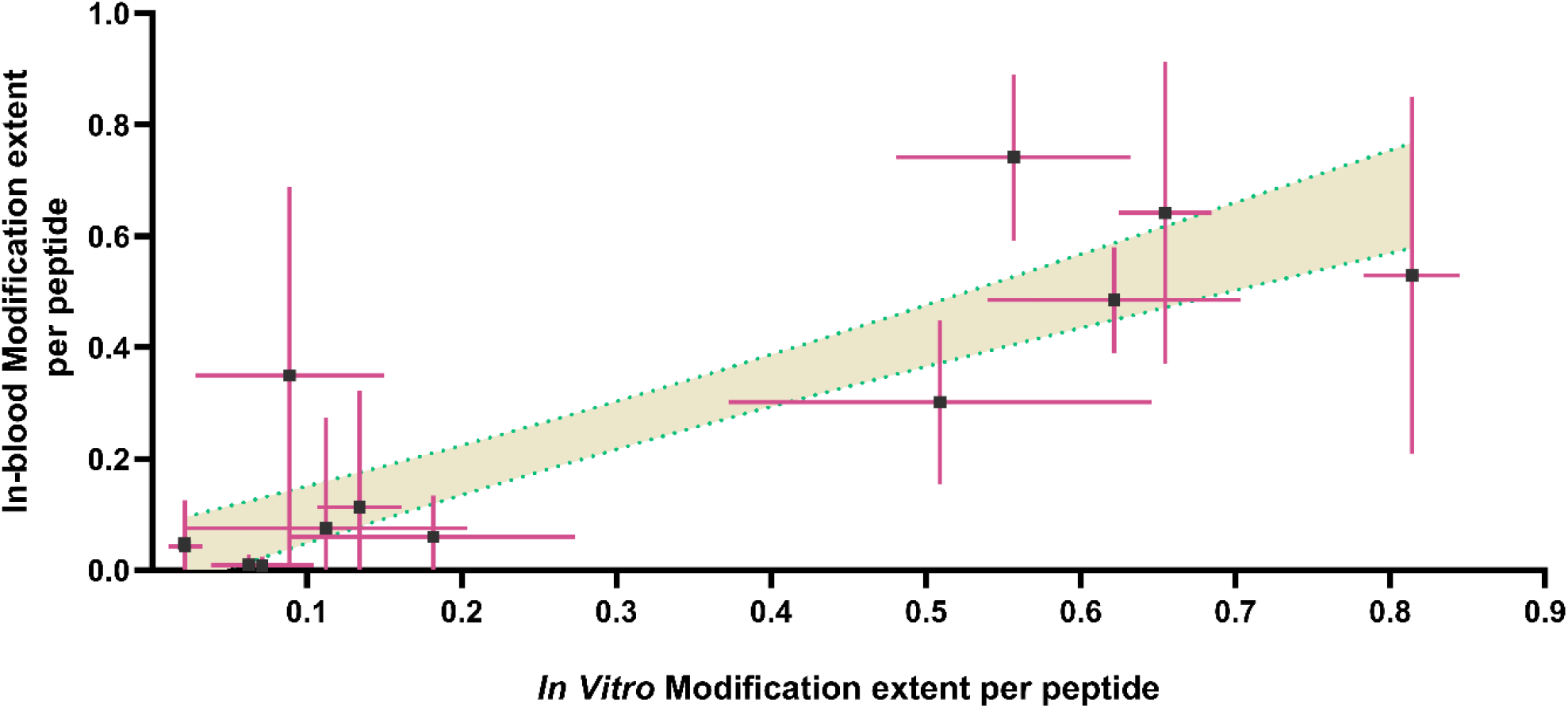
Comparison of *in vitro* and in-blood SO_4_^−^• RPF of protein complement c3. Error bars represent one standard deviation.

## DISCUSSION

A driving motivation for this study was to address unmet technological needs in the structural characterization of biotherapeutics and disease-related proteins in complex clinical matrices. T2DM represents a prime example of a condition where structural proteomics could yield transformative insights. T2DM is characterized by complex systemic changes involving chronic inflammation, insulin resistance, and metabolic dysfunction—processes in which protein conformation and post-translational modifications play critical roles^28, 29, 30^. Complement c3 has emerged as a significant contributor but also a potential effector of disease progression of T2DM^31, 32, 33, 34^. Elevated complement c3 levels are associated with insulin resistance, chronic low-grade inflammation, and metabolic dysfunction—hallmarks of T2DM^31, 32, 35^. Its activation products, such as complement c3a, can impair insulin signaling and beta-cell function, suggesting a direct role in metabolic dysregulation^33^. In-blood RPF found significant differences in complement c3 topography in T2DM mice compared to WT, indicating that we have detected a structural basis behind the role of complement c3 in T2DM. Persulfate-based RPF offers a powerful means to probe conformational changes in proteins like complement c3 directly within blood, enabling the study of their structural states in both healthy and diabetic conditions. Such structural insights could reveal hidden aspects of disease biology, leading to the identification of early biomarkers and novel therapeutic targets. Beyond T2DM, in-blood RPF holds potential for applications in a wide range of diseases where protein structure and interactions are altered. Neurodegenerative diseases, autoimmune conditions, and cancer are all characterized by protein misfolding, aggregation, or abnormal interaction networks^36, 37, 38, 39, 40, 41^—phenomena that can be effectively studied through *in situ* structural proteomics.

Future technical enhancements, including improved intracellular labeling, integration with other biofluids, and development of an automated RPF workflow, will broaden the method’s applicability and reproducibility. Work is ongoing to enhance the ability of sodium persulfate to label intra-cellular proteins, enabling the investigation of integral and intracellular analytes. The method’s compatibility with other biofluids such as cerebrospinal and amniotic fluid further broadens its clinical relevance. To facilitate broader implementation, development is underway for an automated RPF workflow. Automation will reduce variability and improve reproducibility, both of which are essential for clinical adoption. The use of the FOX Photolysis platform further streamlines the implementation of this method, supporting its potential for widespread adoption in clinical and translational research settings.

In-blood RPF holds significant potential for both pre-clinical and clinical applications by enabling direct structural analysis of proteins within complex biological matrices. This approach offers a powerful means to identify conformational biomarkers of disease, revealing structural changes in key proteins that may underlie or reflect pathological processes. It is well-suited for use with clinical samples, such as blood, and can be extended to pre-clinical animal models to track disease progression or therapeutic response. Additionally, in-blood RPF provides valuable insights into the structural stability, aggregation, and degradation of protein pharmaceuticals under near-physiological conditions, supporting structural pharmacology efforts in drug development.

## CONCLUSION

In-blood RPF represents a significant advancement in structural proteomics, enabling detailed analysis of protein conformation and interaction in complex clinical matrices such as whole blood. Its application to disease-relevant proteins in T2DM underscores its potential to provide novel mechanistic insights and identify novel early biomarkers and therapeutic targets.

## Supplementary Information

Supplementary Figure S1. Effectiveness of quench solutions; Supplementary Figure S2. Comparison of in-blood RPF of intracellular and extracellular proteins detected in WT and T2DM whole blood; Supplementary Figure S3. Hierarchical clustering heatmaps on the abundance of top intracellular and extracellular proteins detected from T2DM and WT whole blood; Supplementary Table S1. Body weight and blood glucose levels of mice; Supplementary Table S2. Sequence coverage of top 11 proteins in T2DM and WT whole blood.

## ACKNOWLEDGMENT

The authors would like to thank GenNext Technologies for providing the FOX Footprinting system. The authors gratefully acknowledge Home Place Pastures (Como, MS) for generously providing bovine blood, Dr. Harry Fyke (University of Mississippi) for the generous contribution of canine blood, and Dr. Michael Bouvet (University of California San Diego School of Medicine) for providing initial wild-type mouse blood used in method development. This research was funded by NIH grant R01GM127267. L.M.J. acknowledges funding from NIH grant R35GM144324. This research was supported by the Analytical and Biophysical Chemistry Core and the Imaging Core of the Glycoscience Center of Research Excellence, which is supported by an Institutional Development Award (IDeA) from the National Institute of General Medical Sciences of the National Institutes of Health under award number P20GM130460.

## References

1. Wayment-Steele HK, et al. Predicting multiple conformations via sequence clustering and AlphaFold2. Nature 625, 832–839 (2024).

2. Chance MR. Unfolding of apomyoglobin examined by synchrotron footprinting. Biochem Biophys Res Commun 287, 614–621 (2001).

3. Kaur P, Kiselar J, Yang S, Chance MR. Quantitative protein topography analysis and high-resolution structure prediction using hydroxyl radical labeling and tandem-ion mass spectrometry (MS). Mol Cell Proteomics 14, 1159–1168 (2015).

4. Xie B, Sood A, Woods RJ, Sharp JS. Quantitative Protein Topography Measurements by High Resolution Hydroxyl Radical Protein Footprinting Enable Accurate Molecular Model Selection. Sci Rep 7, 4552 (2017).

5. Chance MR, Farquhar ER, Yang S, Lodowski DT, Kiselar J. Protein Footprinting: Auxiliary Engine to Power the Structural Biology Revolution. J Mol Biol 432, 2973–2984 (2020).

6. Olson LJ, et al. Allosteric regulation of lysosomal enzyme recognition by the cation-independent mannose 6-phosphate receptor. Commun Biol 3, 498 (2020).

7. Wang K, et al. Cryo-EM reveals the architecture of placental malaria VAR2CSA and provides molecular insight into chondroitin sulfate binding. Nat Commun 12, 2956 (2021).

8. Li X, et al. Structural Analysis of the Glycosylated Intact HIV-1 gp120-b12 Antibody Complex Using Hydroxyl Radical Protein Footprinting. Biochemistry 56, 957–970 (2017).

9. Cheng Z, Misra SK, Shami A, Sharp JS. Structural Analysis of Phosphorylation Proteoforms in a Dynamic Heterogeneous System Using Flash Oxidation Coupled In-Line with Ion Exchange Chromatography. Anal Chem 94, 18017–18024 (2022).

10. Shortt RL, Wang Y, Hummon AB, Jones LM. Development of Spheroid-FPOP: An In-Cell Protein Footprinting Method for 3D Tumor Spheroids. J Am Soc Mass Spectrom 34, 417–425 (2023).

11. Espino JA, Jones LM. In Vivo Hydroxyl Radical Protein Footprinting for the Study of Protein Interactions in Caenorhabditis elegans. J Vis Exp, (2020).

12. Gau BC, Chen H, Zhang Y, Gross ML. Sulfate radical anion as a new reagent for fast photochemical oxidation of proteins. Anal Chem 82, 7821–7827 (2010).

13. McKenzie-Coe AA, Johnson DT, Peacock RB, Zhang Z, Jones LM. Evaluating the Sulfate Radical Anion as a New Reagent for In-Cell Fast Photochemical Oxidation of Proteins. Journal of the American Society for Mass Spectrometry 32, 1644–1647 (2021).

14. Sharp JS, et al. Flash Oxidation (FOX) System: A Novel Laser-Free Fast Photochemical Oxidation Protein Footprinting Platform. J Am Soc Mass Spectrom 32, 1601–1609 (2021).

15. Iwase T, et al. A Simple Assay for Measuring Catalase Activity: A Visual Approach. Scientific Reports 3, 3081 (2013).

16. Xie B, Sharp JS. Hydroxyl Radical Dosimetry for High Flux Hydroxyl Radical Protein Footprinting Applications Using a Simple Optical Detection Method. Anal Chem 87, 10719–10723 (2015).

17. Espino JA, Mali VS, Jones LM. In Cell Footprinting Coupled with Mass Spectrometry for the Structural Analysis of Proteins in Live Cells. Anal Chem 87, 7971–7978 (2015).

18. Aebi H. Catalase. In: Methods of Enzymatic Analysis (Second Edition) (ed Bergmeyer HU). Academic Press (1974).

19. Roush AE, Riaz M, Misra SK, Weinberger SR, Sharp JS. Intrinsic Buffer Hydroxyl Radical Dosimetry Using Tris(hydroxymethyl)aminomethane. J Am Soc Mass Spectrom 31, 169–172 (2020).

20. Misra SK, Orlando R, Weinberger SR, Sharp JS. Compensated Hydroxyl Radical Protein Footprinting Measures Buffer and Excipient Effects on Conformation and Aggregation in an Adalimumab Biosimilar. AAPS J 21, 87 (2019).

21. Misra SK, Sharp JS. Enabling Real-Time Compensation in Fast Photochemical Oxidations of Proteins for the Determination of Protein Topography Changes. J Vis Exp, (2020).

22. Sharp JS, Misra SK, Persoff JJ, Egan RW, Weinberger SR. Real Time Normalization of Fast Photochemical Oxidation of Proteins Experiments by Inline Adenine Radical Dosimetry. Anal Chem 90, 12625–12630 (2018).

23. Zhang B, Cheng M, Rempel D, Gross ML. Implementing fast photochemical oxidation of proteins (FPOP) as a footprinting approach to solve diverse problems in structural biology. Methods (San Diego, Calif 144, 94–103 (2018).

24. Shami AA, Misra SK, Jones LM, Sharp JS. Dimethylthiourea as a Quencher in Hydroxyl Radical Protein Footprinting Experiments. J Am Soc Mass Spectrom 34, 2864–2867 (2023).

25. Espino JA, Zhang Z, Jones LM. Chemical Penetration Enhancers Increase Hydrogen Peroxide Uptake in C. elegans for In Vivo Fast Photochemical Oxidation of Proteins. J Proteome Res 19, 3708–3715 (2020).

26. Espino JA, Jones LM. Illuminating Biological Interactions with in Vivo Protein Footprinting. Anal Chem 91, 6577–6584 (2019).

27. Johnson DT, Punshon-Smith B, Espino JA, Gershenson A, Jones LM. Implementing In-Cell Fast Photochemical Oxidation of Proteins in a Platform Incubator with a Movable XY Stage. Anal Chem 92, 1691–1696 (2020).

28. Gonzalez-Dominguez A, et al. Catalase post-translational modifications as key targets in the control of erythrocyte redox homeostasis in children with obesity and insulin resistance. Free Radic Biol Med 191, 40–47 (2022).

29. Kojta I, Chacinska M, Blachnio-Zabielska A. Obesity, Bioactive Lipids, and Adipose Tissue Inflammation in Insulin Resistance. Nutrients 12, (2020).

30. Su T, et al. Myeloid-derived grancalcin instigates obesity-induced insulin resistance and metabolic inflammation in male mice. Nat Commun 15, 97 (2024).

31. King RJ, et al. Fibrinogen interaction with complement C3: a potential therapeutic target to reduce thrombosis risk. Haematologica 106, 1616–1623 (2021).

32. Nishimura T, et al. Clinical significance of serum complement factor 3 in patients with type 2 diabetes mellitus. Diabetes Res Clin Pract 127, 132–139 (2017).

33. Pereira de Lima R, et al. C3aR1 on beta cells enhances beta cell function and survival to maintain glucose homeostasis. Mol Metab 96, 102134 (2025).

34. Ajjan RA, Schroeder V. Role of complement in diabetes. Mol Immunol 114, 270–277 (2019).

35. Ghosh P, Sahoo R, Vaidya A, Chorev M, Halperin JA. Role of complement and complement regulatory proteins in the complications of diabetes. Endocr Rev 36, 272–288 (2015).

36. Chiti F, Dobson CM. Protein Misfolding, Amyloid Formation, and Human Disease: A Summary of Progress Over the Last Decade. Annu Rev Biochem 86, 27–68 (2017).

37. Garcia-Alonso L, Holland CH, Ibrahim MM, Turei D, Saez-Rodriguez J. Benchmark and integration of resources for the estimation of human transcription factor activities. Genome Res 29, 1363–1375 (2019).

38. Guimaraes LE, Baker B, Perricone C, Shoenfeld Y. Vaccines, adjuvants and autoimmunity. Pharmacol Res 100, 190–209 (2015).

39. Liu Y, Beyer A, Aebersold R. On the Dependency of Cellular Protein Levels on mRNA Abundance. Cell 165, 535–550 (2016).

40. Shahnawaz M, et al. Discriminating alpha-synuclein strains in Parkinson’s disease and multiple system atrophy. Nature 578, 273–277 (2020).

41. Soto C, Pritzkow S. Protein misfolding, aggregation, and conformational strains in neurodegenerative diseases. Nat Neurosci 21, 1332–1340 (2018).

