## Supplementary Information for "Radical Footprinting in Mammalian Whole Blood"

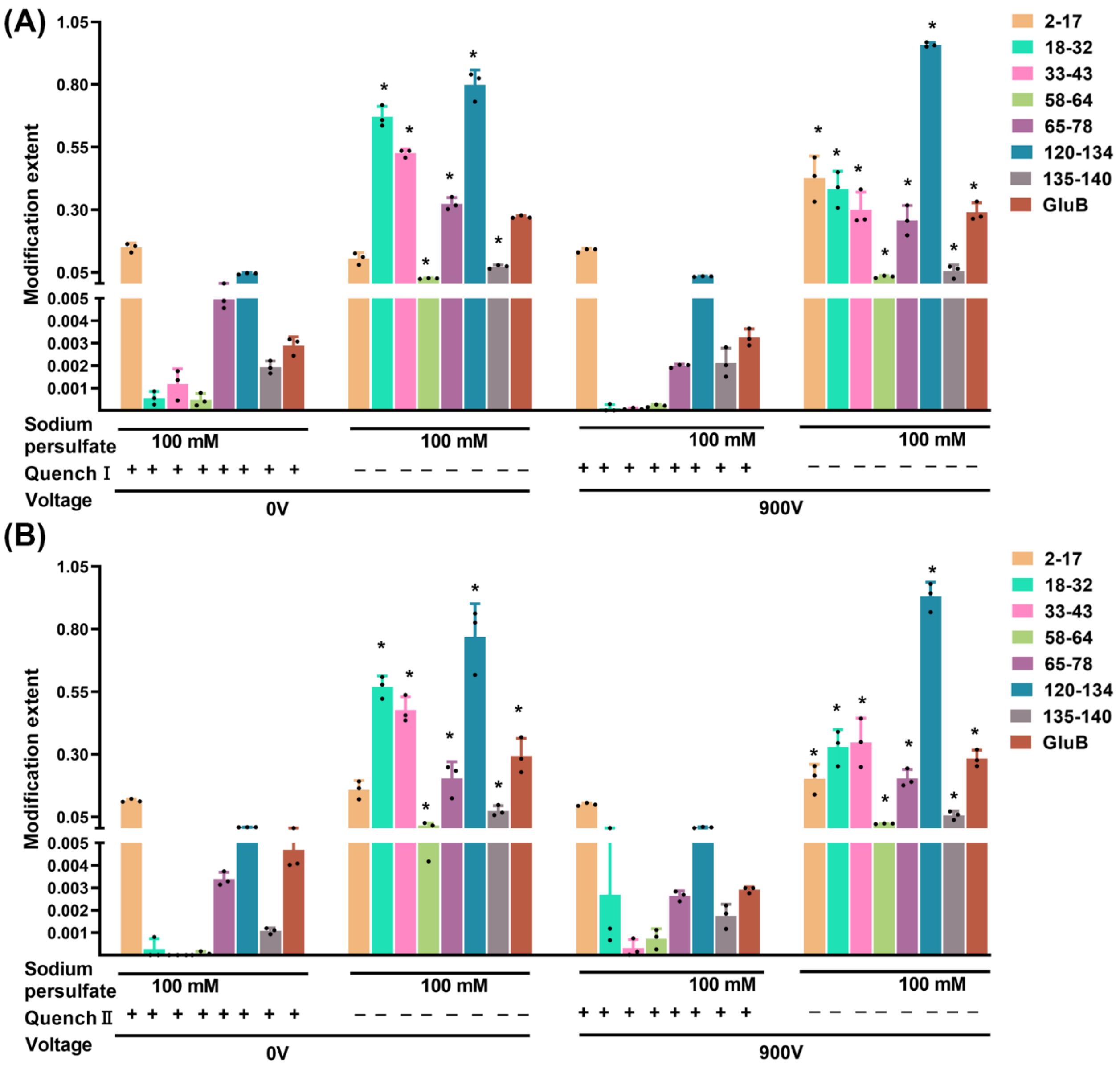


**Figure S1. Effectiveness of quench solutions. (A) Evaluation of Quench I (without imidazole); (B) Evaluation of Quench II (with imidazole).** Both quenches were found to significantly reduce oxidation of most peptides with or without radical photoproduction. Statistical significance was calculated by a two-tailed unpaired t-test ($\alpha = 0.05$) versus the quenched control under the same conditions; peptides showing a significant difference in oxidation are marked with an asterisk. Myoglobin peptides are denoted by their position. The myoglobin peptide corresponding to the 2–17 residues displays significant in-source oxidation. Error bars represent one standard deviation.

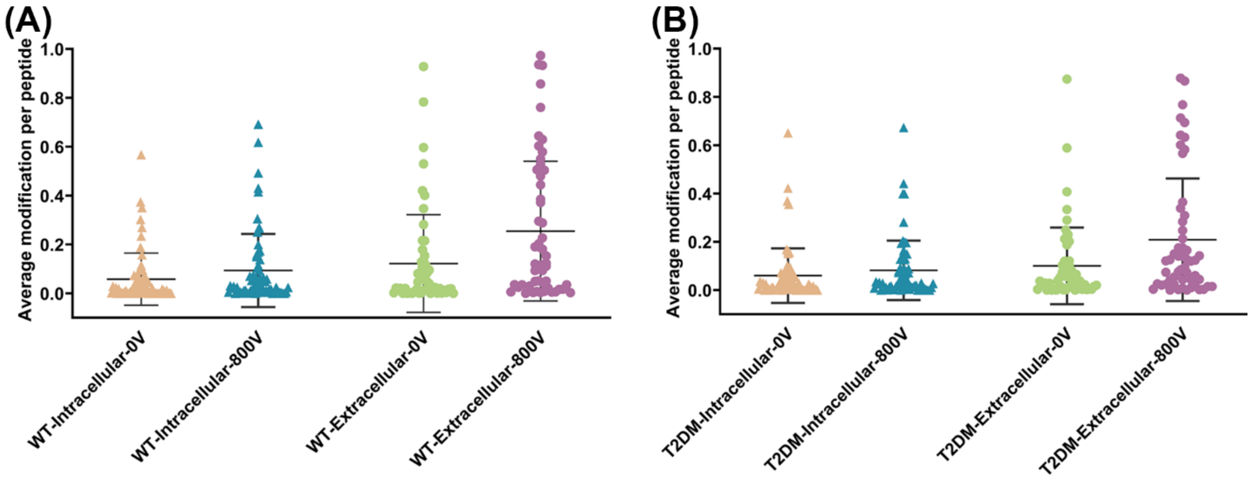


**Figure S2. Comparison of in-blood RPF of intracellular and extracellular proteins detected in WT and T2DM whole blood. (A)** Comparison of modification extent of intracellular and extracellular proteins of WT. Each data point represents a detected peptide; **(B)** Comparison of modification extent of intracellular and extracellular proteins of T2DM. Each data point represents a detected peptide.

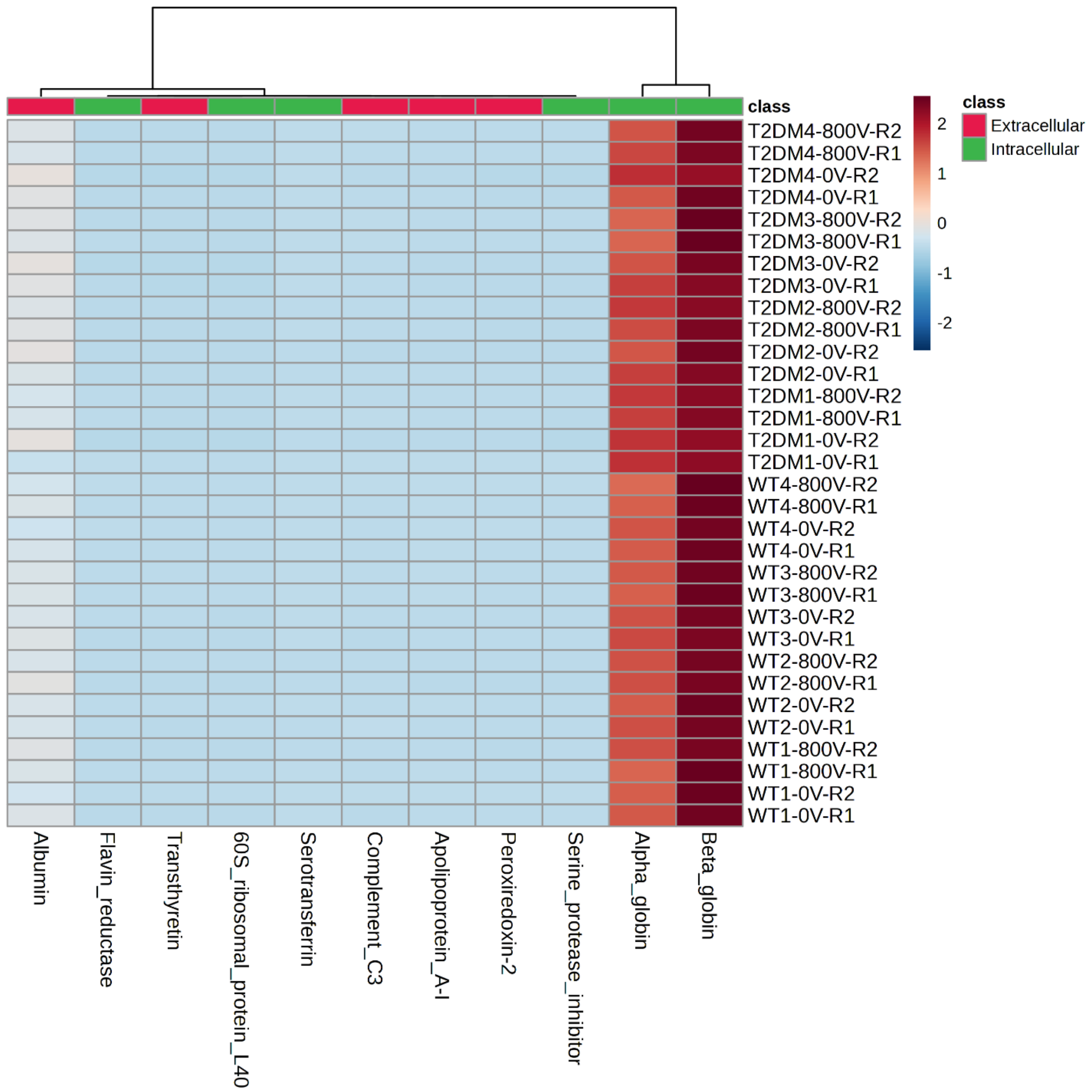


**Figure S3. Hierarchical clustering heatmaps on the abundance of top intracellular and extracellular proteins detected from T2DM and WT whole blood.** Each colored cell on the map corresponds to an intensity value. Red: high; blue: low.

**Table S1. Body weight and blood glucose levels of mice**

| Mice | Boday Weight (mg) | Body Glucose(mg/dL) |
|---|---|---|
| T2DM-1 | 49.09 | 500 |
| T2DM-2 | 47.04 | 586 |
| T2DM-3 | 48.38 | 546 |
| T2DM-4 | 45.64 | 586 |
| WT-1 | 32.53 | 156 |
| WT-2 | 34.90 | 172 |
| WT-3 | 40.13 | 293 |
| WT-4 | 39.12 | 185 |

**Tabel S2. Sequence coverage of top 11 proteins in T2DM and WT whole blood.**

| Protein Name | Average sequence coverage (%) | | Class |
|---|---|---|---|
| | T2DM (n=4) | WT (n=4) | |
| Alpha globin | 99.84 ± 0.29 | 99.95± 0.18 | Intracellular |
| Beta globin | 99.48 ± 0.82 | 99.83 ± 0.30 | Intracellular |
| Flavin reductase | 75.49 ± 5.48 | 82.25 ± 6.28 | Intracellular |
| Transthyretin | 76.34 ± 8.38 | 78.54 ± 9.09 | Extracellular |
| Peroxiredoxin-2 | 75.99 ± 4.66 | 75.22 ± 1.21 | Intracellular |
| Apolipoprotein A-I | 73.39 ± 7.78 | 73.44 ± 7.89 | Extracellular |
| Complement c3 | 70.59 ± 4.35 | 73.04 ± 4.64 | Extracellular |
| Serotransferrin | 70.35 ± 4.21 | 70.45 ± 4.97 | Extracellular |
| Albumin | 72.15 ± 2.89 | 70.29 ± 4.07 | Extracellular |
| Serine protease inhibitor | 60.69 ± 4.82 | 60.49 ± 5.40 | Intracellular |
| 60S ribosomal protein L40 | 52.06 ± 1.84 | 52.05 ± 2.10 | Intracellular |
